# Metabolomics characterisation of cassava pre-breeding populations with enhanced whitefly tolerance

**DOI:** 10.1101/2025.02.05.636665

**Authors:** L Perez-Fons, A Bohorquez-Chaux, MI Gomez-Jimenez, LA Becerra Lopez-Lavalle, PD Fraser

## Abstract

Cassava (*Manihot esculenta* Crantz) provides food security for over 500 million people in Sub-Saharan Africa (SSA). Whitefly (*Bemisia tabaci*) is a pest in this region that result in ca. 50% crop yield losses. Thus, it is important to develop approaches that will generate new varieties tolerant to this pest to advance food security in the region. Two parental cassava varieties, ECU72 tolerant to whiteflies and COL2246 a susceptible line have been used to generate bi-parental populations. The F1 generation has been screened for whitefly resistance and progeny identified displaying enhanced tolerance. From designated F1 tolerant progeny, F2 families have been generated and phenotyped. The tolerance to whiteflies in the F2 population was further enhanced. Untargeted metabolomics was used to characterise whitefly susceptible and tolerant sub-groups. PCA of the molecular features generated clustering of accessions into whitefly resistant and susceptible groups and differentiating metabolite biomarkers were identified. The most significant metabolite marker for resistance being the chemical feature 316.0924. Although not consistent among all whitefly resistance sub-groups targeted LC-MS analysis revealed several pathways displaying perturbed levels. These include cyanogenic glycosides, apocarotenoids and phenylpropanoid super-pathway comprising of hydroxycinnamic acids, flavonoids and proanthocyanidins. Thus, the generation of a bi-parental population for whitefly tolerance/susceptibility enabled the identification of quantitative metabolite markers, the pathways contributing to tolerance, the underlying modes of action associated with resistance and the potential for the development of future high-throughput low-cost proxy markers. The approach also provides generic insights into future breeding strategies utilising bi-parental progeny for the enhancement of traits.

**SUMMARY.**

**SIGNIFICANCE STATEMENT.**

## INTRODUCTION

Cassava (*Manihot esculenta Crantz*) is a staple crop for many low and middle income countries (LMICs) populations. In Africa alone over 500 million people are dependent on cassava and its products for food security. Its ability to withstand drought conditions and poor soils make it a potential climate resilient crop (Parmar et al., 2017). Given these key socio-economic criteria significant investments have been made to advance cassava breeding as a means of addressing food/nutritional security. Genomic based approaches including marker assisted selection (MAS) and genomic selection (GS) are the current approaches for accelerating cassava breeding programs in Sub-Saharan Africa (SSA) (Okogbenin et al., 2007, Okogbenin et al., 2012, Wolfe et al., 2017, Esuma et al., 2022, Rabbi et al., 2022). These GS models and marker discovery studies have focused on exploiting SSA genetic resources and using phenotypic measurements mostly related to biomass and plant vigour (Wolfe et al., 2016, Wolfe et al., 2017, Wolfe et al., 2021). The diversity of the SSA cassava genetic pool used in breeding programs is limited to multiple genetic crossings of existing germplasm (Reinhardt Howeler, 2013, Rabbi et al., 2022, Ferguson et al., 2023, Xia et al., 2023), with occasional introduction of diverse Latin American germplasm (Okogbenin et al., 2007, Okogbenin et al., 2012).

Considering the level of investment in automation and computational power available to modernise cassava breeding programs in SSA countries, comparatively little attention has been taken to address the foundational resources and tools that underpin traditional breeding programs, namely genetic diversity, accurate phenotype determinations and robust trait discovery pipelines (Alemu et al., 2024). Although genetic tools directed to breeder implementation are prioritised, examples of discovery-based outputs augmenting breeding strategies are now becoming evident. For example, the Cassava Source-Sink (CASS) project, providing a comprehensive understanding of the metabolic constraints and trade-offs between dry matter and provitamin A traits in roots, as well as changes in the above-ground vegetative biomass (Sonnewald et al., 2020, Gutschker et al., 2024). Tolerance to whitefly infestation has also been facilitated by the characterisation of Latin American diverse germplasm, while metabolomic and transcriptomic analysis has led to potential mechanism of action being proposed (Perez-Fons et al., 2019, Irigoyen et al., 2020, Nye et al., 2023). Infestation of cassava crops by whiteflies (*Bemisia tabaci*) can result in crop yield losses of up to 50% (Sam et al., 2024) and is thus a major target for cassava breeding. In the present work, phenotypic and metabolome characterisation of bi-parental populations bred for whitefly resistance in cassava are presented. The aim of the study was to (i) use metabolite profiling as a complementary phenotypic tool for the selection of progeny, (ii) identify phenotypic markers at metabolome level, (iii) identify underlying modes of action and (iv) indicate the use of these populations as pre-breeding genetic resources.

## RESULTS

### Genetic resources and phenotypic classification

Two cassava accessions ECU72 and COL2246, with contrasting whitefly phenotypes, resistant and susceptible respectively, have been characterised at metabolome and transcriptome level and a mechanism of action proposed (Perez-Fons et al., 2019, Irigoyen et al., 2020, Nye et al., 2023). Both varieties were used to generate a number of bi-parental crosses in order to: (i) assess phenotype heritability, (ii) obtain pre-breeding material with stable and transferrable whitefly resistance phenotype, and (iii) traceable phenotype markers.

A first generation (F1) of bi-parental populations was established by crossing female flowers of the whitefly resistant (WF-R) variety ECU72 and male flowers of susceptible (WF-S) varieties COL2246 and TMS60444 from Latin American and African germplasm backgrounds respectively. This strategy was defined by the characteristic male flower sterility of ECU72. The resulting F1 families, designated as CM8996 and GM8586 according to CIAT’s internal nomenclature system, was then evaluated for whitefly phenotypes under multiple growing conditions using the automated nymph count system Nymphstar (Bohorquez-Chaux et al., 2023). Both families presented a number of siblings showing enhanced WF-R than that of the parent ECU72 (Fig. 1A, B) when screened over multiple seasons, soil conditions and propagation methods. The average phenotypic determinations for each family compared to the parents shows CM8996 being closer to WF-R parent, and GM8586 being closer to the WF-S parent (Supplementary Fig. S2). Segregating phenotypes were evident in both cases with CM8996 presenting extremer whitefly phenotypes. Percentiles 15 and 85 of nymph counts were chosen as cut-off points to designate extreme WF-R and WF-S phenotype classes respectively. The difference between means of WF-R and WF-S classes in CM8996 family was 1430 nymphs whilst in GM8586 the difference was 1026 (Supplementary Fig. S1, Supplementary File S01). Additionally, the mean difference between WF-R or WF-S classes and the intermediary phenotype (WF-I) was 662 and 768 nymphs in CM8996, but 440 and 586 in GM8586. In all cases, the differences between phenotype classes were highly significant (p<0.0001).

**Figure 1:**
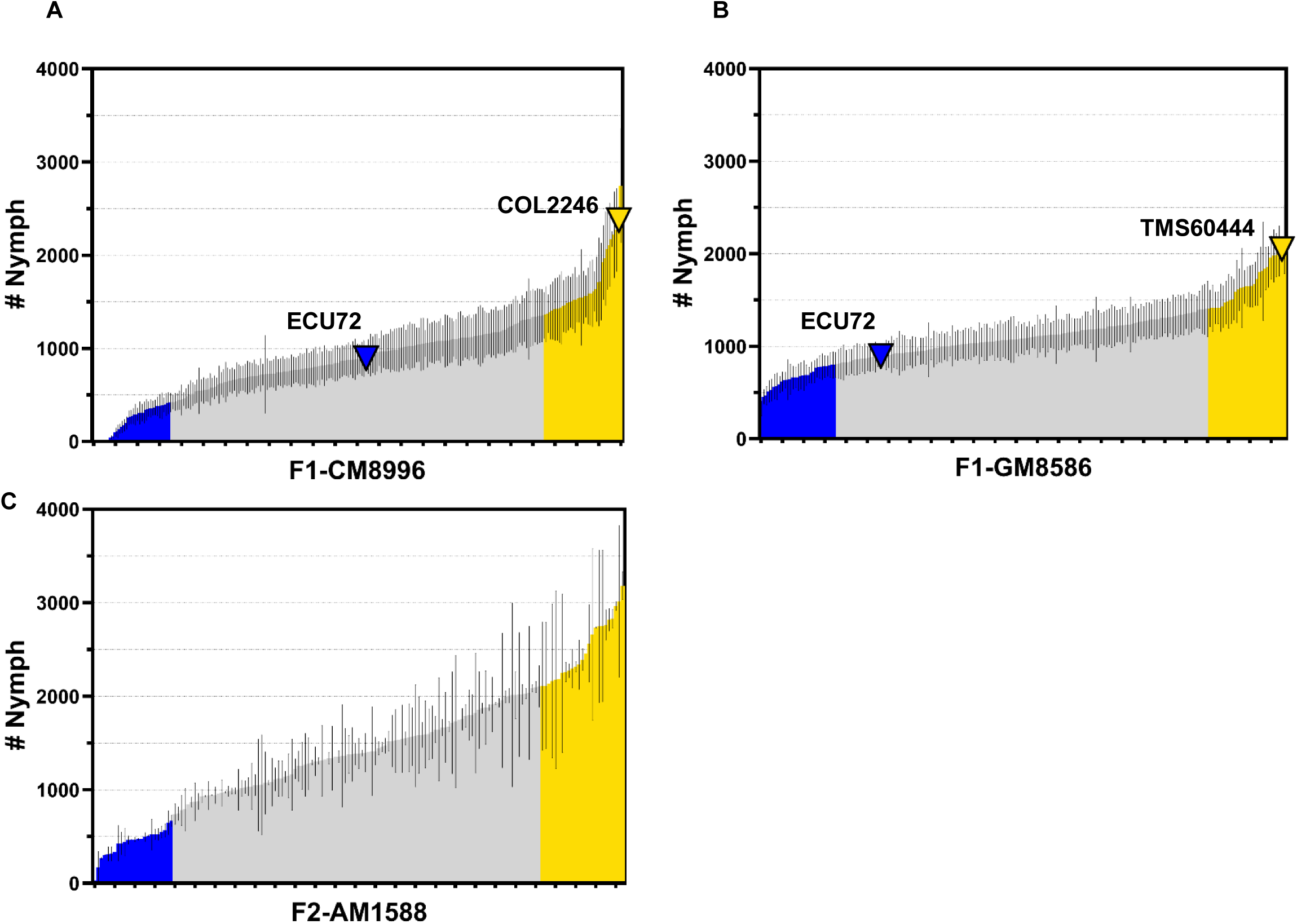
Phenotyping of populations as whitefly nymph counts of F1 families CM8996 (**A**) and GM8586 (**B**), and the F2 population AM1588 (**C**). Percentiles 15 and 85 are coloured as extreme WF-R (blue) and WF-S (yellow) phenotypic groups, respectively. Parents are indicated with inverted triangle symbols and siblings as bars with standard error of mean of the different experiments. Siblings with intermediary phenotype are coloured in light grey.

Subsequently, advancement into second generation of crosses (F2) were performed by crossing siblings with highest WF-R in CM8996 in order to produce pre-breeding plant material with stable/fixed WF-R phenotype and reduced heterozygosity. Despite the male flower sterility of female parent ECU72, sibling CM8996-199 displayed enhanced flowering with viable male flowers thus enabling a true-F2 population by selfing CM8996-199.

Phenotypic classification of true-F2 AM1588 (Fig. 1C) indicates that extreme WF-R class differs significantly (p<0.0001) from both WF-I and WF-S classes by 978 and 2068 nymph counts respectively, and a mean’s difference of 1090 nymphs separates significantly WF-S from WF-I classes (Supplementary Fig. S1, Supplementary File S02).

### Metabolome overview of bi-parental crosses

An untargeted acquisition and data analysis approach was used for evaluating metabolome variation of bi-parental populations F1 and F2 of ECU72. The number of chemical features extracted from the LC-MS analysis of each family was 3285 in CM8996, 4278 in GM8586, and 2284 in AM1588 (Supplementary files S03, S05, S07). After filtering outliers and inconsistent missing values, the final untargeted data matrix used for subsequent analysis contained 2899 variables for CM8996, 4252 for GM8586, and 1384 variables for AM1588 (Supplementary files S04, S06, S08).

Components 1 and 2 of principal component analysis (PCA) explain 29.5% and 24.4% variation in the F1 families CM8996 and GM8586 respectively (Fig. 2A, B), and 25.3% in the true-F2 family AM1588 (Fig. 2C). The score plots of both F1s present a wider spread of phenotypic classes WF-R, WF-S and WF-I, whilst F2 population AM1588 shows the extreme phenotypes (WF-R and WF-S) separating from the rest of the siblings presenting intermediary phenotype (Fig. 2C). When extreme phenotype classes WF-R and WF-S were analysed independently by PCA and HCA, clusters within the WF-R and WF-S groups can be withdrawn. These groups were designated as extreme pheno-metabotypes and used for comparative analysis and biomarkers identification.

**Figure 2:**
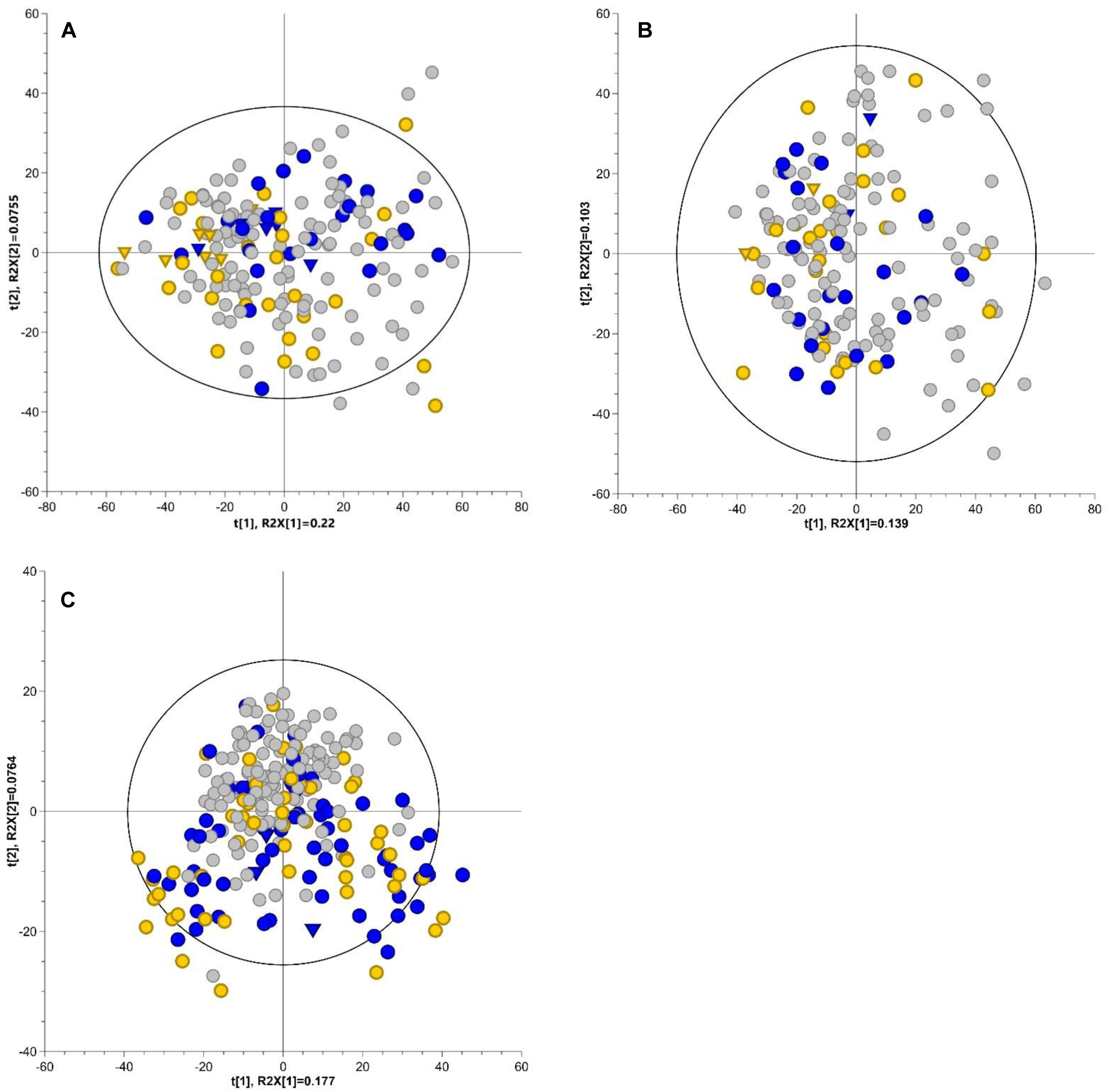
Principal component analysis of populations’ metabolome. Score plot of untargeted LC-MS dataset of F1 family CM8996 (**A**), F1 family GM8586 (**B**) and F2 family AM1588 (**C**). Percentiles 15 and 85 coloured as extreme WF-R (blue) and WF-S (yellow) phenotypic groups. Parents represented with inverted triangle symbols and siblings as circles.

### Metabolite markers of whitefly resistance

A range of supervised machine learning methods are available for biomarker identification in metabolomic studies (Anwardeen et al., 2023, Ghosh et al., 2020). In the present work, orthogonal partial least square discriminant analysis (OPLS-DA) has been used to shortlist candidate metabolites associated with whitefly tolerance. The OPLS-DA model descriptors obtained initially from the analysis of all the siblings selected based on the phenotype classification (15^th^ and 85^th^ quartiles of nymph count) of the F1 family CM8996 presented R2(cum) and Q2(cum) values of 0.484 and 0.0922, respectively, with only one predictive (P1) component and no orthogonal (O) components explaining the variation between and within both phenotypic classes. Similarly, the analysis of the extreme phenotypic classes of the F2 family AM1588 produce a model with R2(cum) fitting value of 0.978 and predictive Q2(cum) value of 0.444. However, the R2X(cum) of the predictive component P1 was substantially lower (0.012) compared to the sum of orthogonal components (0.444). Therefore, to refine/increase the accuracy of the model, the OPLS-DA was performed using groups of extreme pheno-metabotypes, i.e., individuals of each phenotypic class clustering closely and opposite its phenotypic counterpart in the PCA score plot. In the F1-CM8996, one metabotype sub-cluster from each extreme WF-R and WF-S phenotypic class can be withdrawn (Fig. 3A, B). The siblings in the WF-R pheno-metabotype class of CM8996 included 199, 246, 255, 68, 759 and 773, and those in the WF-S pheno-metabotype were 905, 894, 888, 875, 867, 862 and 836. In the F2-AM1588, at least three sub-groups of extreme WF-R pheno-metabotypes, WF-R1, WF-R2 and WF-R3, were observed together with one group of WF-S pheno-metabotype (Fig. 3C, D). The F2 family also included biological replicates of the selected extreme phenotypes, therefore grouping of replicates was an additional selection criterion applied in this case. Those siblings presenting high biological variation, i.e., inconsistent clustering of biological replicates in the PCA score plot were not selected. The list of siblings in the AM1588 WF-S subgroup were 214, 207, 90, 13, 18, 29. The metabotypes included in the different F2’s WF-R sub-groups were 83, 15, 69 and 36 in the WF-R1; 233, 175, 136 and 21 in the WF-R2; 186, 127, 151, 142 and 44 in the WF-R3.

**Figure 3:**
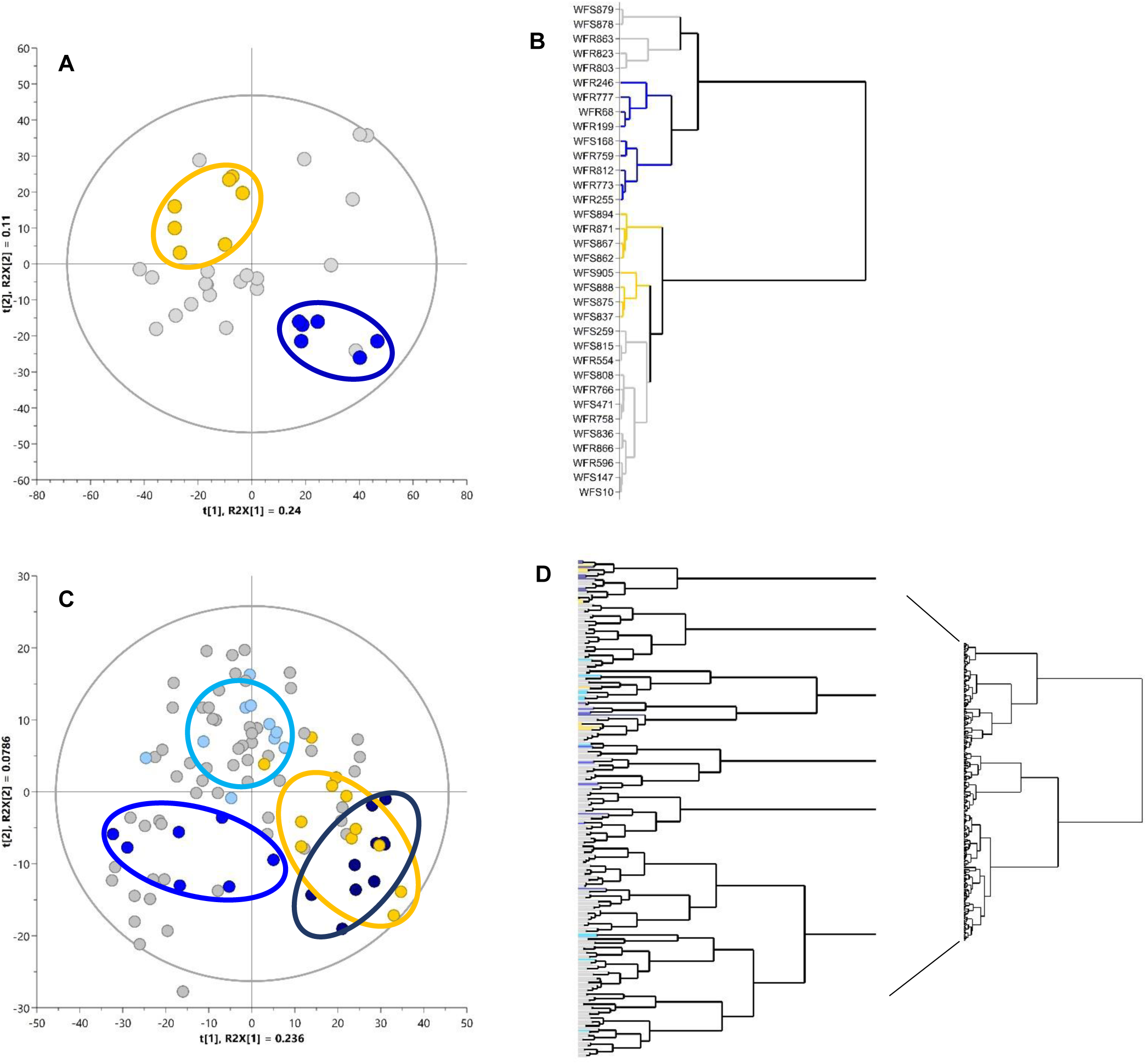
Principal component analysis of selected extreme phenotypes of F1 family CM8996 (**A**) and F2 family AM1588 (**C**). The dendrograms of the populations’ extreme phenotypic groups highlighting the selected pheno-metabotypes of F1 (**C**) and F2 (**D**) and coloured accordingly. The grey dots represent those extreme phenoytpes not selected as pheno-metabotype groups.

Following selection of extreme pheno-metabotypes, an OPLS-DA was applied to pairs of WF-R and WF-S pheno-metabotypes in each bi-parental family F1 and F2 to identify differentiating chemical features that correlate with whitefly resistance or susceptibility phenotype. The data analysis workflow is represented in Fig.4 using F1-CM8996 extreme pheno-metabotypes as an example, and same rationale was applied to the pairs of pheno-metabotypes found in the F2 (Supplementary Figs 3-5). In all cases, the predictive component of the model P1 defining class variation was higher than the orthogonal component O1 (or sum of them) defining the inherent class variation. Moreover, the cumulative R2 and Q2 values of the models representing respectively the level of phenotype variation explained or predicted by metabolome, were over 0.8 [Supplementary File S09].

**Figure 4:**
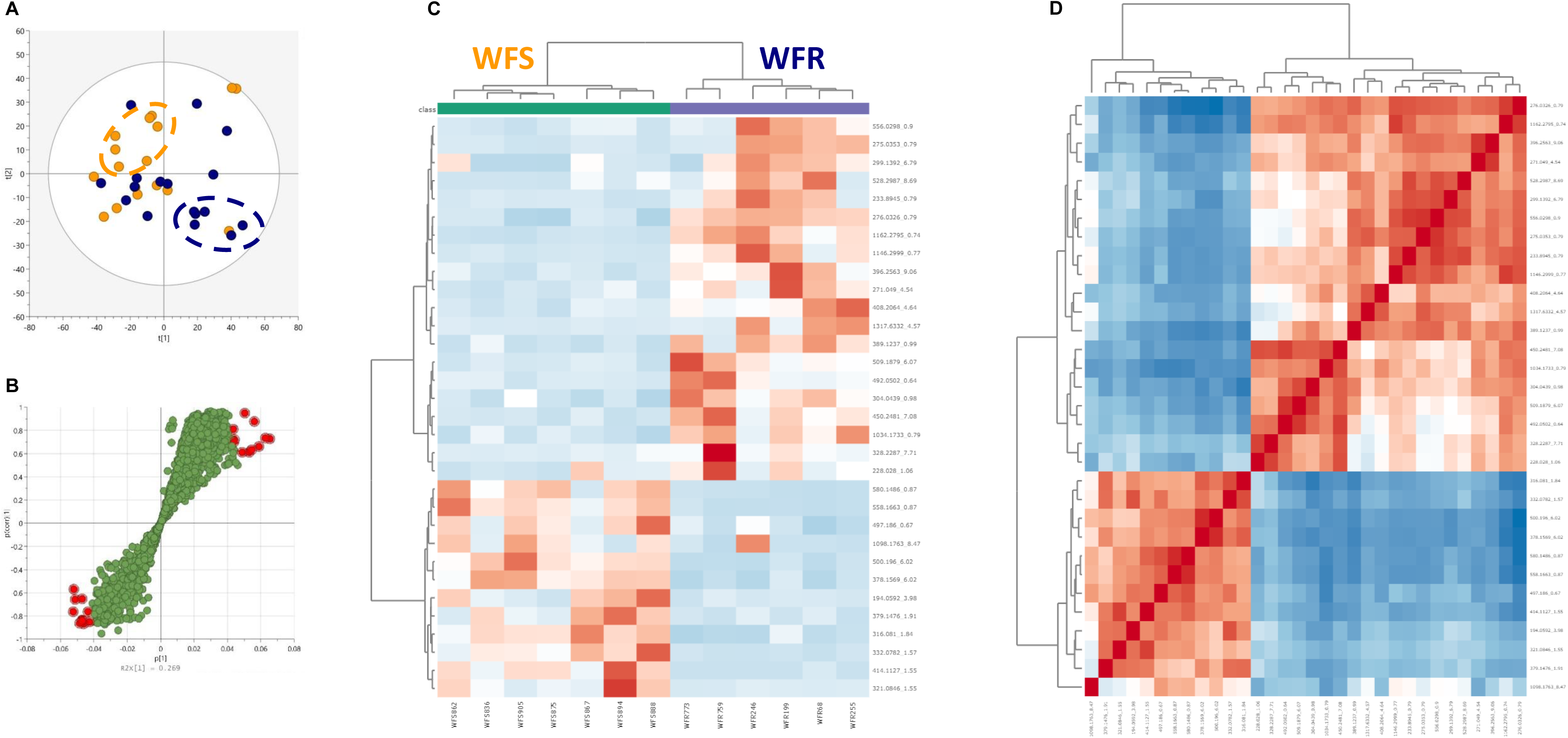
Data analysis workflow followed for the identification of metabolite markers explaining phenotypic differences of F1’s extreme pheno-metabotypes. (**A**) PCA scores plot of F1-CM8996 extreme phenotypic classes, dark blue represent whitefly resistant (WF-R) siblings and yellow dots represent whitefly susceptible (WF-S) siblings. Clusters of pheno-metabotypes selected for OPLS-DA analysis are circled. (**B**) Loadings S-plot obtained using the selected WF-R and WF-S pheno-metabotypes. Red dots indicate those chemical features presenting high correlation and covariance with phenotypic classes and selected as list of markers. (**C**) Heat-map and dendrogram of chemical features retrieved from the S-plot, and (**D**) metabolite-metabolite correlation (Pearson’s) heat-map of selected markers.

The loadings S-plot of each OPLS-DA model was performed to extract a list of candidate markers by selecting those loadings located at the extremes of the S-tails (Fig.4B, Supplementary Figs. 3-5, panel B). In most cases the correlation and covariance cut-off values were 0.6 and 0.04 respectively. Some exceptions include a negative correlation cut-off value of 0.5 in the CM8996 model, or positive cut-off variance of 0.06 in the WF-R2 and WF-R3 models of the F2 family AM1588. A list of 32 chemical features was obtain from the S-plot of CM8996. Similarly, models comparing the different sub-groups of WF-R pheno-metabotypes in the F2 family with the WF-S class provided with lists of 87, 58 or 64 candidates (Supplementary File S10). The heatmaps of the biochemical markers shortlisted showed reciprocal accumulation and correlation between WF-R and WF-S class in each model (Fig. 4C, D; Supplementary Fig.S3-5, panels C, D).

Only one chemical feature, 316.0924_5.34, was identified in all three WF-R subgroups. The WF-R2 and R3 shared the largest number of metabolite markers (13), whilst WF-R1 shares 2 and 4 chemical features with WF-R2 and R3 respectively (Supplementary Fig. S7).

### Targeted metabolite profile of whitefly resistant sub-classes in F2 family AM1588

Using an in-house metabolite library customised for cassava, the untargeted LC-MS data matrix was annotated. The metabolite LC-MS library included a wide range of chemical classes from primary metabolism (mono and disaccharides, amino acids, organic acids or fatty acids) to secondary metabolism (cyanogenic glycosides, hydroxycinnamic acids, hydroxybenzoates, flavonoids superfamily or apocarotenoids), among other miscellaneous compounds (Supplementary file S11). The resulting targeted data matrix contained 169 chemical features annotated (Supplementary file S12) and this was used as input for statistical analysis. Pair-wise statistical comparison of WF-R subgroups and WF-S group of F2 family AM1588 was performed and significant changes of metabolites mapped onto biosynthetic pathways as WF-R/WF-S ratios (Fig. 5). In addition, detailed compositional differences are also summarised as heatmaps in Supplementary File S13.

**Figure 5:**
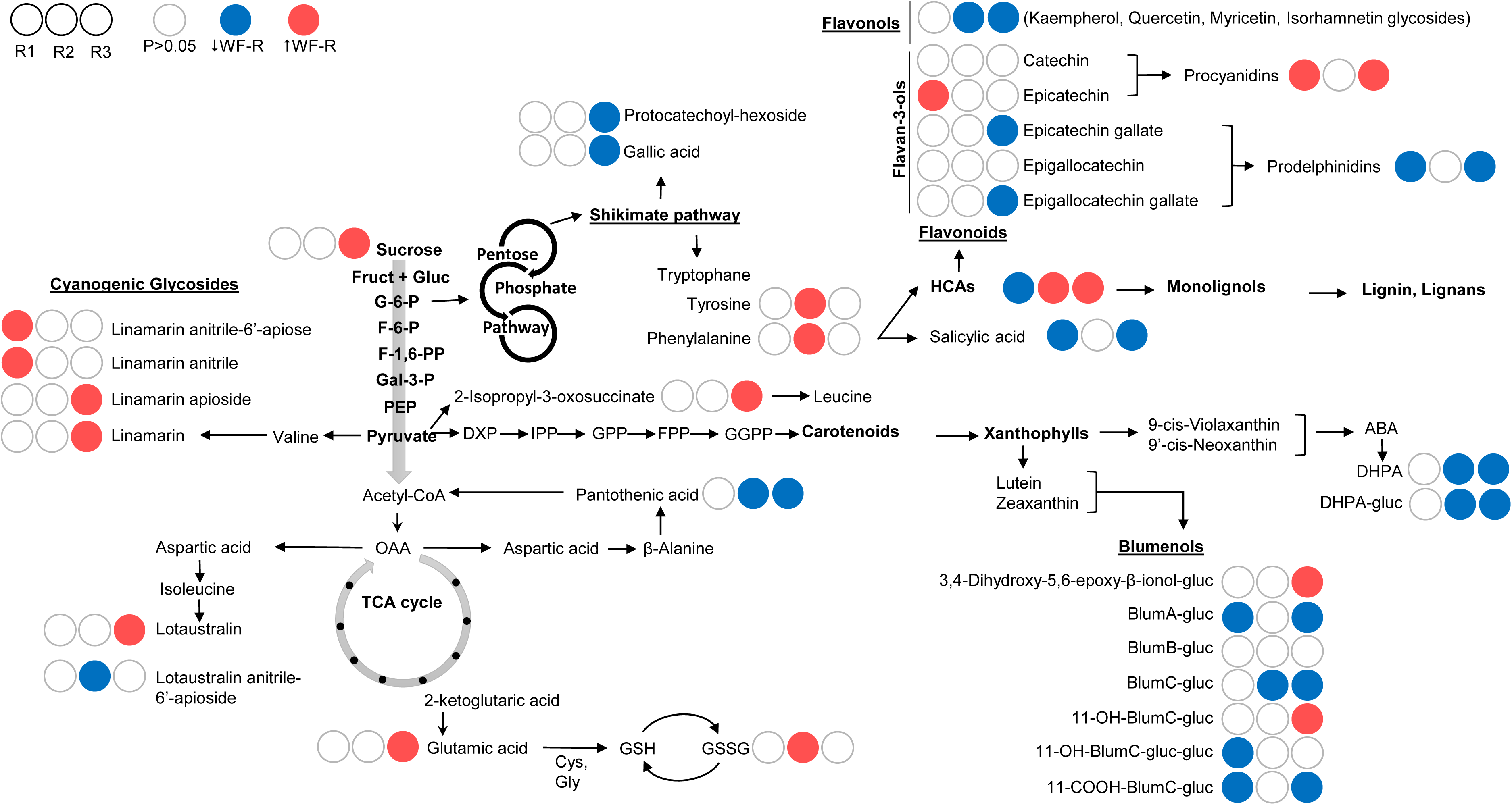
Pathway display illustrating the significant changes between F2’s whitefly resistant sub-groups WF-R1, R2 and R3 compared to the whitefly susceptible (WF-S) sub-group. Red filled circles indicate concentration of metabolite is significantly (p<0.05) higher in the WF-R subgroup, blue coloured circles indicate significantly (p<0.05) lower concentration in the WF-R sub-group than the WF-S group.

The number of significant changes and the identity of the metabolites involved was different across the three WF-R subgroups. The number of significant changes increased with the distance/position between the pheno-metabotypes classes in the PCA plot in the following order: WF-R1 < WF-R2 < WF-R3, with 14 significant metabolites differentiating WF-R1 and WF-S, 25 significant differences between WF-R2 and WF-S and 61 significant differences between WF-R3 and WF-S.

The WF-R1 presented higher levels of anitrile derivatives (non-active) of cyanogenic glycosides, whilst active forms were significantly higher in WF-R3 and no significant changes were detected in the WF-R2 except for the anitrile form of lotaustralin apiose. Similarly, different apocarotenoids profile was observed. All WF-R subgroups display significantly lower concentrations of blumenols in general. The 11-OH-blumenol C glucoside and the 3,4-dihydroxy-5,6-epoxy-beta-ionol glucoside were the only apocarotenoids significantly higher in one of the resistant subgroups (WF-R3). Its oxidised by-product 11-COOH-blumenol C glucoside, the precursor blumenol C glucoside and blumenol A glucoside were statistically lower in WF-R3 and R1 but not in WF-R2. In addition, both catabolites of abscisic acid (ABA), dihydrophaseic acid (DHPA) and its glucoside, were significantly lower in WF-R2 and R3.

The phenylpropanoids super-pathway comprising hydroxycinnamic acids (HCAs) and flavonoids was also significantly different across all WF-R subgroups. Hydroxycinnamic acids involved in the monolignols biosynthesis were statistically higher in both WF-R2 and R3 but lower in the WF-R1 when compared to the susceptible group. In contrast, flavonols and flavan-3-ols were generally lower in WF-R3 and only epicatechin was higher in the WF-R1 sub-group. Polymerised forms of flavan-3-ols, procyanidins and prodelphinidins presented a contrasting trend, with significantly higher concentrations of procyanidins and lower levels of prodelphinidins in WF-R1 and R3 but no statistical changes in WF-R2. The WF-R2 sub-group presented significant reciprocal changes in HCAs and flavonols, as well as in the amino acids phenylalanine and tyrosine, which are the committed precursors of the phenylpropanoids biosynthesis.

## DISCUSSION

Previous studies demonstrate that different sectors of metabolism and a reprogramming of gene expression triggered by a contrasting phytohormones signalling, are involved in the whitefly tolerance of cassava’s natural variation (Perez-Fons et al., 2019, Irigoyen et al., 2020, Nye et al., 2023,). The wide range of molecular and biochemical events suggest an underlying complex genetic trait.

One of the strategies used by breeders to dissect complex genetic traits (QTLs) into minor QTLs that can be effectively transferred to farmers-preferred or elite varieties in a stable manner is to create pre-breeding material via RIL or NIL approaches. This enhances gene dosage of the trait, its markers and reduces epistasis (Scott et al., 2020). These procedures are feasible in crops where botanical seeds are the propagation standards and genetic/biological resources and toolkits are well characterised. That is the case of maize, tomato (Perez-Fons et al., 2014, Tripodi et al., 2021), peppers, rice or cotton, among many others. Cassava is a monoecious plant characterised by separate male and female flowers that mature asynchronously (protogyny) contributing to a high rate of male flowers abortion and limited germination efficiency, challenges that are exacerbated by genetic and environmental factors influencing reproductive processes (Njoku et al., 2015, Bandeira et al., 2021, Long et al., 2024); thus, favouring clonal propagation as the standard method of choice. In addition, the frequency of natural cross-pollination via insect vectors, usually bees, is higher than self-pollination which contributes to enhance the heterozygosity nature of the crop (Alves, 2002, Rodrmguez et al., 2023).

The present work successfully generated and characterised bi-parental populations to develop clones exhibiting enhanced whitefly tolerance or resistance phenotype. These clones also demonstrated reduced heterozygosity and refined genomic regions associated with the trait that could potentially serve as pre-breeding material and true-to-type donors of the whitefly tolerance trait. The populations were characterised both phenotypically and at metabolome level, with the two layers of information integrated and correlated to get deeper insights. The F1s families using ECU72 as trait donor, confirmed heritability of whitefly tolerance across progeny, regardless of the genetic background—Latin American (COL2246) or African (TMS60444). Subsequent selfing crosses produced segregating families, with extreme phenotypes becoming more distinct. These observations suggest increased gene dosage effects or dominance driven by heterosis and epistatic interactions (Birchler et al., 2010, Fujimoto et al., 2018).

It has been reported that cassava’s genetic diversity is correlated to metabolome composition (metabotypes) associated to specific trait/phenotypes (Perez-Fons et al., 2023). Metabotypes—distinct profiles of metabolites within an organism—can provide insights into complex genetic interactions, such as epistasis, that traditional molecular markers might not fully elucidate. By analysing metabolic traits, researchers can uncover interactions between genes that influence phenotypic outcomes, offering a more comprehensive understanding of genetic architecture. For example, a study on *Arabidopsis thaliana* investigated the genetic basis of leaf molybdenum levels and identified significant statistical epistasis between a cis-trans predictor pair affecting this trait. This interaction arose from high-order linkage disequilibrium (LD) among four polymorphisms associated with molybdenum levels, suggesting that metabolic profiling can uncover complex genetic interactions that might be overlooked when relying solely on molecular markers (Zan et al., 2018). Similarly, a study on rice observed transgressive segregation for kilo-grain weight in a recombinant inbred line population, finding that epistasis and complementary gene action adequately accounted for this phenomenon. This supports the idea that metabolic profiles can provide insights into genetic interactions that molecular markers alone might miss (Mao et al., 2011).

In this work, the extreme phenotypes of F1 and F2 populations also present distinctive metabolite profiles (metabotypes). When overlaying extreme phenotypes and metabotypes, only 1 cluster of whitefly resistance siblings can be identified in the F1 CM8996, whilst at least 3 subgroups of metabotypes are identified within the whitefly resistant cluster in the F2 AM1588. The statistical predictive model OPLS-DA indicate that metabolome variation of the selected extreme pheno-metabotypes explains accurately the phenotypic differences, with each phenotypic sub-group presenting a specific metabolite profile composition. This could indicate that the genetic components (small-QTLs) contributing to the complexity of whitefly resistance trait start to segregate independently, thus providing valuable genetic resources for dissecting causal molecular and biochemical mechanisms of the trait, which would ultimately be translated into more accurate markers. The WF-R1 sub-group included the most resistant clones of the F2 family and yet presented the lowest significant differences when compared to the WF-S group, whilst the WF-R2 and R3 sub-groups were markedly different to WF-S group. Despite the quantitative variability in pathways affected, there are common sectors of metabolism perturbed. For example, the phenylpropanoid super-pathway. This supports previous findings (Perez-Fons et al., 2019) that cell wall enhancement through altered precursor flux could be a component of the mechanism. Phytohormones, particularly abscisic acid levels are altered in most of the resistant sub-groups. This finding is in agreement with previous reports (Irigoyen et al., 2020, Nye et al., 2023). Finally, it is interesting that some blumenols are also affected as this opens new lines of investigation linked to the potential contribution/involvement by arbuscular mycorrhizal fungi (AMF) and stress tolerance (Wang et al., 2018).

Detailed analysis of the allelic variation among the genomic regions of interest will provide a selection of candidate genes that could refine the region further and contribute to the underlying molecular and biochemical mechanisms underlying the trait of interest. Incorporating the metabolomics and other -omics at early stages of breeding programs would also support accuracy of genomic selection models by assisting the assembly and cross-validation process of the training populations, typically used to build genomic prediction models and design breeding population. The approach would ultimately reduce phenotyping costs and improve model accuracy. In the present study whitefly tolerance has been used as the target trait, but the findings have generic implications that can benefit the current cassava breeding pipelines established in SSA through projects such as NEXTGEN (https://www.nextgencassava.org/) or RTB-Breeding (https://rtbbreeding.cgiar.org/). For example, whereby introgression of a donor F1 or F2 line with an enhanced phenotype into the recipient background (Fig. 6). The trait will be stronger, presumably due to increased zygosity but the potential to capture epistasis and heterosis is retained. In addition, the phenotypic strength will alleviate the inherited segregation from the heterozygosity of the recipient line. Synergistic to the approach are the potential for molecular and quantitative biochemical markers as well as systems level analysis for deciphering complex modes of action.

**Figure 6:**
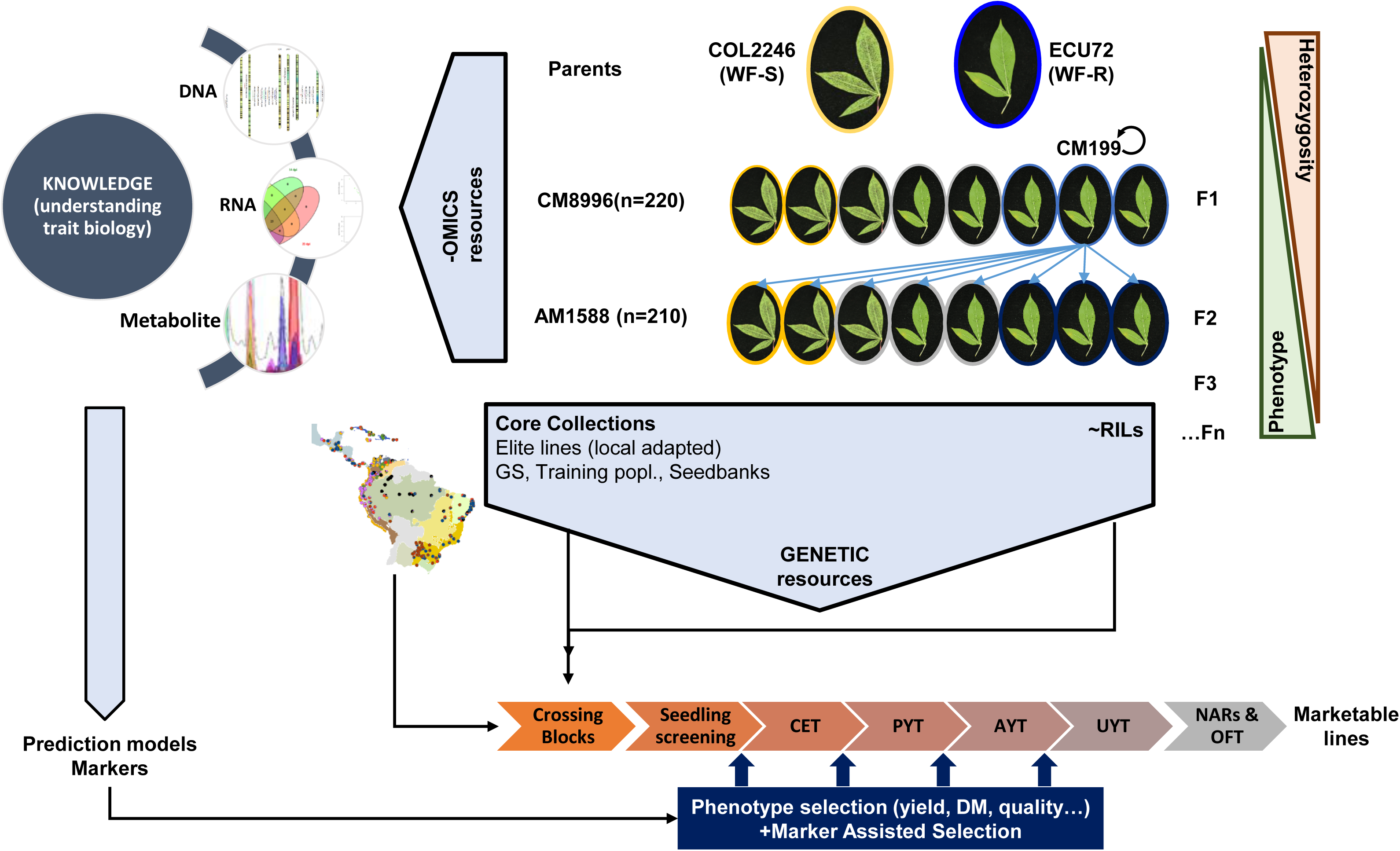
Schematic diagram of alternative breeding strategy integrating discovery/knowledge outputs into current cassava breeding pipelines. CET, clonal evaluation trials; PYT, preliminary yield trials; AYT, advanced yield trials; UYT, uninform field trials; OFT, on-farm trials; NARs, national agronomic research institutions; DM, dry matter; GS, genomic selection; RILs, recombinant inbred lines.

With the advent of AI and machine-learning algorithms, metabolomics capacity to predict phenotype accurately is becoming closer to implementation in plant breeding (Fernie and Alseekh, 2022). Although the cost and high throughput metabolomics is still not comparable to sequencing technologies, the examples reported up to date suggest that models’ accuracy built from metabolome data outperform those based on genomic information (Colantonio et al., 2022).

In summary, the present study has furthered valuable tools and resources for breeding new cassava accessions for whitefly tolerance. The approach used has contributed to elucidating underlying mechanisms associated with whitefly tolerance and has generic implications for added value to current cassava breeding programmes.

## METHODS

### Plant growing conditions

All bioassays were made at the CIAT experimental station. Shoot tips from in vitro-grown cassava genotypes were excised and grown in 17 N rooting medium for 30 days in the cassava tissue culture lab at CIAT. These included whitefly susceptible genotypes (parentals COL2246 and TMS60444; check COL1468), one whitefly-resistant genotype (parental ECU72), 220 progenies of the F_1_ CM8996, 198 progenies of the F_1_ GM8586, and 210 progenies of the F_2_ AM1588.

For hardening, the plants were sown in 2-L plastic bags with sterile soil, in a ratio of 1:3 sand to black soil (no clay topsoil). Plants were grown in a glasshouse with temperatures ranging from 24 to 28 °C under a long-day light cycle (16-h light/ 8-h dark).

### Mass rearing of *Aleurotrachelus socialis* and whitefly bioassays

The *A. socialis* colony was raised on *Manihot esculenta* var. COL1468 as previously described by (Bohorquez-Chaux et al., 2023). For the whitefly-infestation bioassays, each two-month-old cassava genotypes were put into a mesh cage of 18 m Large × 3 m Wide × 3 m High to confine the whitefly adults after infestation in a glasshouse. Infestations were initiated by releasing ∼22,000 adults of *A. socialis* into cages. When the adult whiteflies were removed at 3 dpi, the two youngest infested leaves, which are preferred by whiteflies for feeding and egg deposition, were tagged for future collection.

Three biological replicates were used for each genotype. Infested plants were placed in a randomized complete block design (RCBD). To capture the effect of each life stage of the whitefly on the cassava plants, the sample collection time points were chosen to represent landmarks during the *A. socialis* life cycle. Leaf tissues were harvested prior to infestation (time zero), 1 day post-infestation (1 dpi; adult feeding and egg deposition), and 14 dpi (2nd and 3rd instar feeding). After collection, leaves were frozen in liquid nitrogen and stored at −80 °C. Subsequently, leaves collected were freeze-dried (Christ LyoCube®) for 2 days and then shipped at ambient temperature in silica gel to analytical facilities at Royal Holloway University of London. Transit period did not exceed 3 days and samples were stored at −80 °C after arrival.

### Phenotyping

For the phenotyping of F1s and F2s and their respective parents and controls, a free-choice experiment was carried out under greenhouse conditions, and an automated counting method called Nymphstar (Bohorquez-Chaux et al., 2023) was used. With this approach, photographs of the infested leaves were analysed using the plugin and data on nymph counts (NC) and area infested by nymphs were obtained.

In order to validate consistency of whitefly resistance phenotype, the infestation trials were performed under different growing condition using plant material from in vitro culture or from stakes and/or micro-stakes transferred from the field. Generally, in vitro grown material was planted onto poor soil (2:1 ratio of black soil: sand), and stakes were planted onto rich soil (3:1, black soil: sand ratio). The experiments performed over different seasons were as follows:

| Year | Cross | Propagation | Substrate |
| --- | --- | --- | --- |
| 2013 | CM8996 | In vitro | poor |
| 2016 | CM8996 | Stake | poor |
| 2016 | GM8586 | Micro-stake | poor |
| 2017 | CM8996 | In vitro | rich |
| 2017 | CM8996 | Stake | rich |
| 2017 | GM8586 | stake | rich |
| 2018 | CM8996 | In vitro | poor |
| 2018 | GM8586 | In vitro | poor |
| 2020 | AM1588 | In vitro | poor |
| 2021 | AM1588 | stake | rich |

### Metabolites extraction

Freeze-dried tissue was ground to fine powder using Qiagen TissueRuptor as described in (Rosado-Souza et al., 2019) and 10mg was used for extraction of metabolites as described in (Perez-Fons et al., 2019). Briefly, 700 μl of 50% v/v methanol was added and the mixture shaken for 1 hr at room temperature. Addition of chloroform (700 μl) followed by centrifugation (3 min, 14000 rpm) allowed separation of semi-polar and non-polar compounds into the epiphase and organic phase respectively. Semi-polar extract was filtered with 0.45 μm nylon membranes and non-polar extract dried under vacuum. Both extracts were kept at −20° C until analysis.

### Untargeted metabolomics analysis by LC-MS

An aliquot of 95 μl of the semi-polar extract (epiphase) was transferred to glass vials and spiked with 5 μl of internal standard (genistein at 0.2 mg/ml in methanol). Samples were kept at 8° C during analysis and volume injection was 1 μl.

For the analysis of the semi-polar extracts a C18 reverse phase column and a UHPLC-ESI-Q-TOF system from Agilent Technologies was used. The analytical platform consisted of a 1290 Infinity II liquid chromatograph, a 6560 Ion mobility Q-TOF mass spectrometer operating in Q-TOF mode only and equipped with an Agilent Jet Stream (AJS) electrospray source. Data was acquired in MS mode from 100 to 1700 mDa under negative electrospray ionisation. Nebulizer and sheath gas temperatures were 325° and 275° C respectively; flowrate of drying and sheath gas (nitrogen) were 5 and 12 L/min respectively and nebulizer pressure 35 psi. Capillary VCap, nozzle and fragmentor voltages were set up at 4000, 500 and 400 V. A reference mass solution was continuously infused to ensure mass accuracy calibration. Compounds were separated in a Zorbax RRHD Eclipse Plus C18 2.1×50 mm, 1.8 μm and 2.5% acetonitrile in water and acetonitrile as mobile phase A and B respectively, both solvents containing formic acid (0.03% vol.) as additive. Gradient started at 2% B for 1 min, increase to 30% B over 5 min, stay isocratic for 1 min followed by an increase to 90% B in two minutes. The washing step was maintained isocratic for two minutes and initial conditions were restored for re-equilibration for 3 minutes. Flowrate and column temperature were set at 0.3 ml/min and 30° C, respectively.

### Processing of untargeted metabolomics data files

Retention time alignment and extraction of chemical features of raw data files were processed with Agilent’s Profinder (version 10.0) software using the Batch Recursive Molecular Feature Extraction mode. The following settings were selected to extract molecular features within a retention time (RT) range of 0.3 to 10 min: peak height threshold 1000 counts, RT tolerance +/- 0.2 min, mass tolerance 10 ppm, chlorine and formic acid adducts, and water neutral losses were also considered. Only molecular features (MF) with matching scores higher than 70 and present in at least 15% of each sample group (QC and samples) were included in the final data matrix. This resulted in the detection of over 2000 MFE per population’s data set. Putative characterisation of chemical identity was inferred from accurate mass values calculated from mass-to-charge ratio (m/z) signals. Chemical formulae were generated using the following elemental constrains: C: 70; H: 140; O: 40; N: 10; S: 5; P: 3 ((Ma et al., 2014), (https://pmn.plantcyc.org/CASSAVA/search-query?type=COMPOUND&formula=C)), formic acid and chlorine adducts and/or multiply charged species (z=1, 2) were also considered. Those chemical formula (up to 5) with the highest score (based on mass difference and isotopic pattern fitting) were selected for blasting against ChemSpider and Dictionary of Natural Products chemical databases. Additionally, an in-house library of cassava metabolites based on chromatographic parameters (retention time) and fragmentation pattern (MS/MS) (Perez-Fons et al., 2019, Perez-Fons et al., 2023) was used to complement and validate findings of the putative identification pipeline described above.

### Statistical analysis and data analysis workflow

Batch correction of LC-MS untargeted data (extracted ion chromatogram peak areas) was applied using quality control samples (QC). In addition, normalisation against area of internal standard was performed to correct for instrument variation. Missing values were input by using the median value of each mass reported from the extraction chemical feature pipeline, and those presenting over 75% of missing values were excluded from analysis. The resulting data matrix was then used as input for multivariate analysis (PCA, HCA and OPLS-DA) in SIMCA-P v17 (Sartorius AG, Germany) and univariate analysis (t-tests, ANOVA, Pearson’s correlation) in Prism v10 (GraphPad software LLC). Centering and pareto-scaling were applied for multivariate analysis. Pair-wise comparisons were performed by multiple two-sample *t-test* assuming unpaired data, Gaussian distribution (parametric) and inconsistent standard deviation (Welch test). Multiple t-test comparisons were corrected with Holm-Sidak post-hoc test setting a threshold value for significance (alpha) at 0.05. Random Forest analysis was performed using MetaboAnalyst v5.0.

## Data Statement

All relevant data supporting results presented are provided in supplemental files. Metabolomics data sets of raw (areas) and processed data (normalized), and phenotyping datasets are accessible in Mendeley Data: Perez-Fons, Laura; Bohorquez-Chaux, Adriana; Fraser, Paul; Becerra Lopez-Lavalle, Luis Augusto (2025), “ACWP-Cassava Mapping Populations Metabolomics”, Mendeley Data, V2, doi: 10.17632/pdbjznbf2g.2

## Acknowledgements

This work was supported by the African Cassava Whitefly Project (ACWP)-Phase 2 funded by Natural Resources Institute (NRI), University of Greenwich, UK, from a grant provided by the Bill and Melinda Gates Foundation (OPP1200124). We thank Mr. Chris Gerrish for technical support, and CIAT’s bioinformatics and entomology team for assistance in the design and conduction of whitefly tolerance phenotyping experiments. We also thank Dr. Winnie Gemode, current CIAT’s cassava geneticist, for reviewing the manuscript.

## Supporting information legends

Supplementary Fig.S1: significant differences of phenotypic classes. ANOVA unpaired Brown-Forsythe tests corrected for unequal variance (Welch’s correction).

Supplementary Fig.S2: Heterosis effect. Bars represent mean and standard deviation of nymph counts. ANOVA unpaired Brown-Forsythe tests corrected for unequal variance (Welch’s correction).

Supplementary Fig.S3: Metabolite markers explaining phenotypic classification of F2’s WF-R1 subgroup and WF-S sub-group.

Supplementary Fig.S4: Metabolite markers explaining phenotypic classification F2’s WF-R2 subgroup and WF-S sub-group.

Supplementary Fig.S5: Metabolite markers explaining phenotypic classification F2’s WF-R3 subgroup and WF-S sub-group.

Supplementary Fig.S6: Venn diagram of metabolite markers identified within the WF-R class of the F2 family AM1588.

Supplementary File S01: Nymph count of F1 families CM8996 and GM8586.

Supplementary File S02: Nymph count of F2 family AM1588.

Supplementary File S03: Untargeted chemical features extracted from the LC-MS analysis of F1 CM8996. Raw data output obtained from Profinder v 10.0.

Supplementary File S04: Untargeted chemical features extracted from the LC-MS analysis of F1 CM8996. Processed data after QC correction and internal standard normalisation and used as input for multivariate and statistical analysis.

Supplementary File S05: Untargeted chemical features extracted from the LC-MS analysis of F1 GM8586. Raw data output obtained from Profinder v 10.0.

Supplementary File S06: Untargeted chemical features extracted from the LC-MS analysis of F1 GM8586. Processed data after QC correction and internal standard normalisation and used as input for multivariate and statistical analysis.

Supplementary File S07: Untargeted chemical features extracted from the LC-MS analysis of F2 AM1588. Raw data output obtained from Profinder v 10.0.

Supplementary File S08: Untargeted chemical features extracted from the LC-MS analysis of F2 AM1588. Processed data after QC correction and internal standard normalisation and used as input for multivariate and statistical analysis.

Supplementary File S09: Descriptors of OPLS-DA models.

Supplementary File S10: List of markers obtained from the loadings S-plot.

Supplementary File S11: Library of metabolite compounds used for annotation of chemical features.

Supplementary File S12: Targeted dataset of F2-AM1588 including annotated compounds and relative quantification.

Supplementary File S13: Targeted dataset of F2-AM1588 including p-values (T-test) and fold-changes (ratios) using WF-S group as reference.

