## Supplemental figures for "Metabolomics characterisation of cassava pre-breeding populations with enhanced whitefly tolerance"

Supplementary Figures

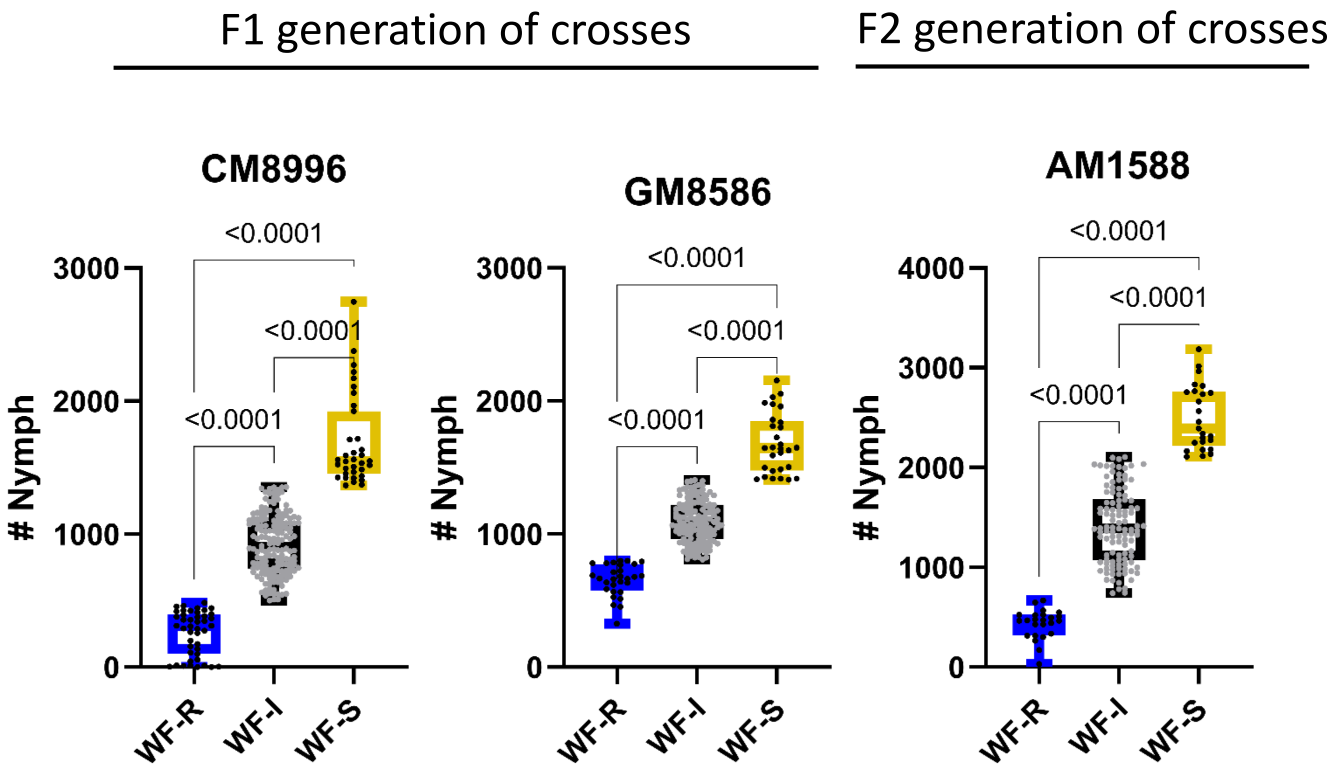

**Supplementary Fig.S1:** significant differences of phenotypic classes. ANOVA unpaired Brown-Forsythe tests corrected for unequal variance (Welch's correction).

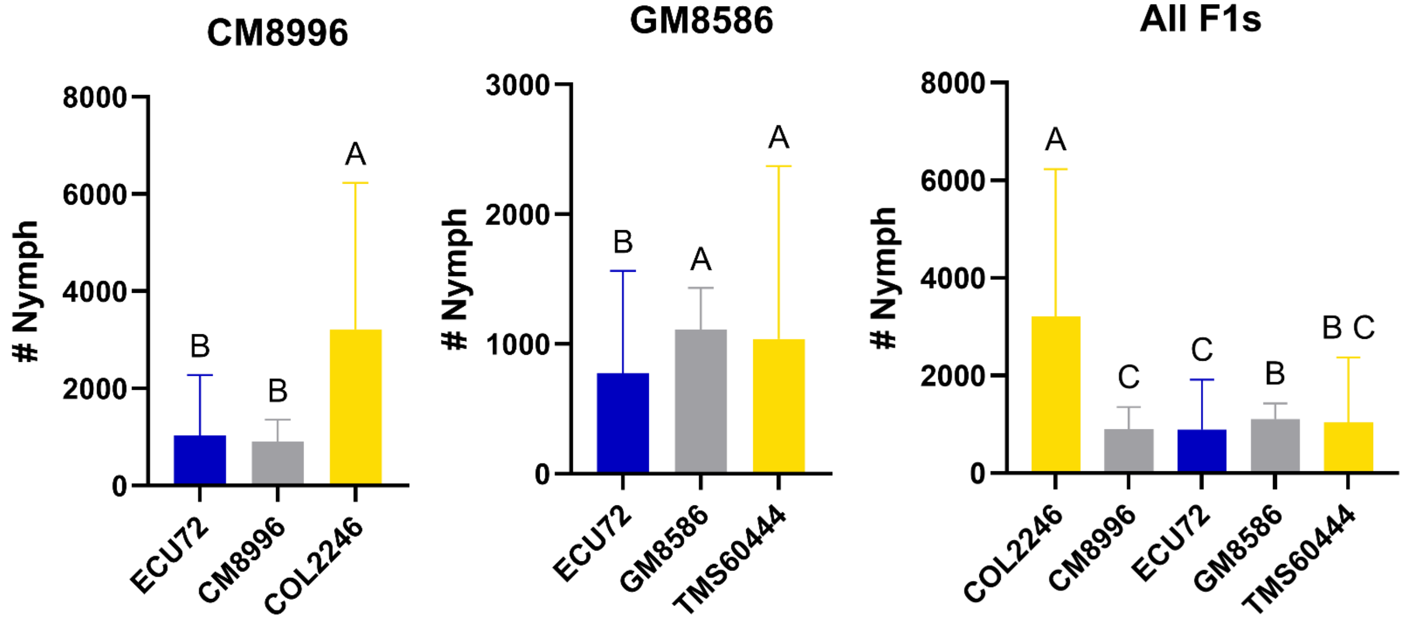

| Table Analysed | CM8996 | GM8586 | All F1s |
|---|---|---|---|
| Brown-Forsythe ANOVA test | | | |
| F* (DFn, DFd) | 103.7 (2.000, 297.1) | 8.543 (2.000, 220.3) | 94.52 (4.000, 356.9) |
| P value | <0.0001 | 0.0003 | <0.0001 |
| P value summary | **** | *** | **** |
| Significant diff. among means (P < 0.05)? | Yes | Yes | Yes |
| Welch's ANOVA test | | | |
| W (DFn, DFd) | 64.02 (2.000, 350.4) | 21.88 (2.000, 287.5) | 39.20 (4.000, 498.3) |
| P value | <0.0001 | <0.0001 | <0.0001 |
| P value summary | **** | **** | **** |
| Significant diff. among means (P < 0.05)? | Yes | Yes | Yes |
| Data summary | | | |
| Number of treatments (columns) | 3 | 3 | 5 |
| Number of values (total) | 706 | 635 | 1341 |

**Supplementary Fig.S2:** Heterosis effect. Bars represent mean and standard deviation of nymph counts. ANOVA unpaired Brown-Forsythe tests corrected for unequal variance (Welch's correction).

### Metabolite markers of extreme metabo/phenotypes

**AM1588_WF-R1**

Heatmap & dendrogram

Metabolite-metabolite correlation (Pearson's)

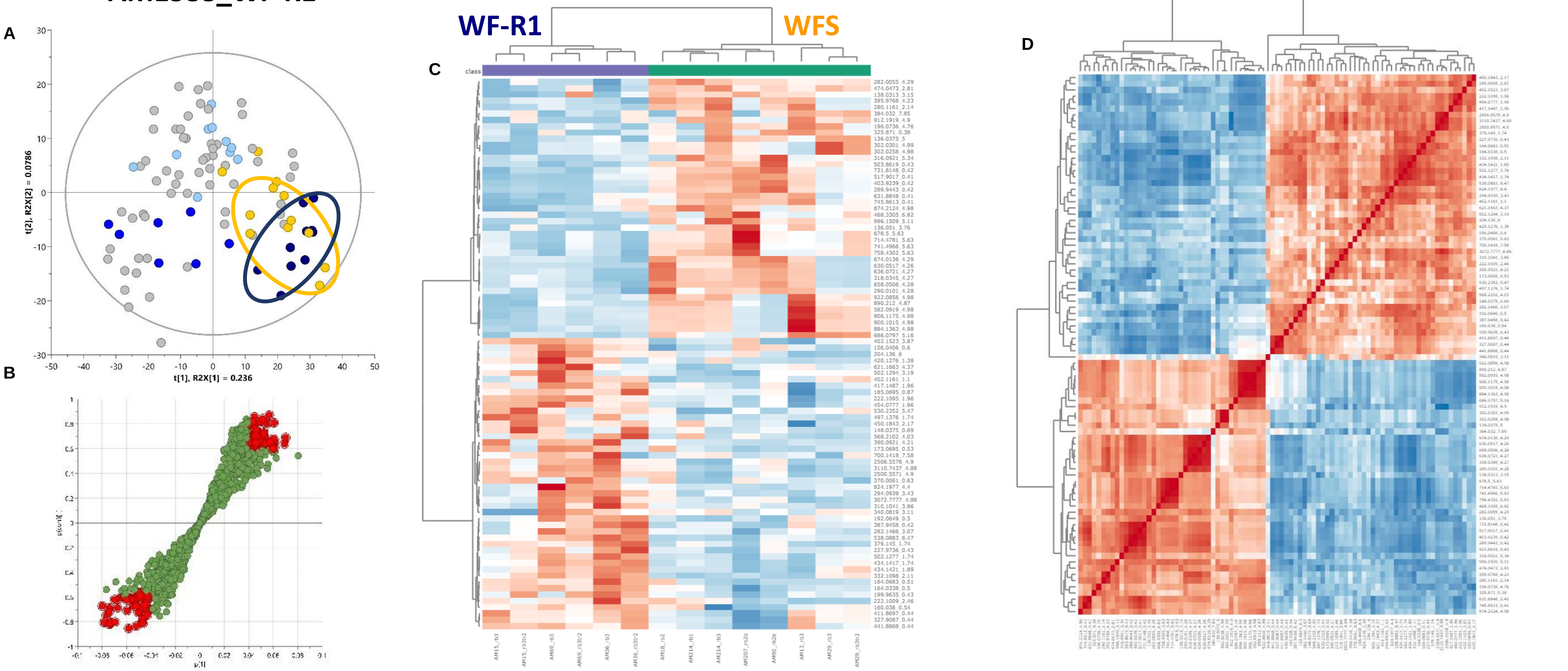

**Supplementary Fig.S3**: Metabolite markers explaining phenotypic classification of F2's WF-R1 subgroup and WF-S group.

### Metabolite markers of extreme metabo/phenotypes

Heatmap & dendrogram

Metabolite-metabolite correlation (Pearson's)

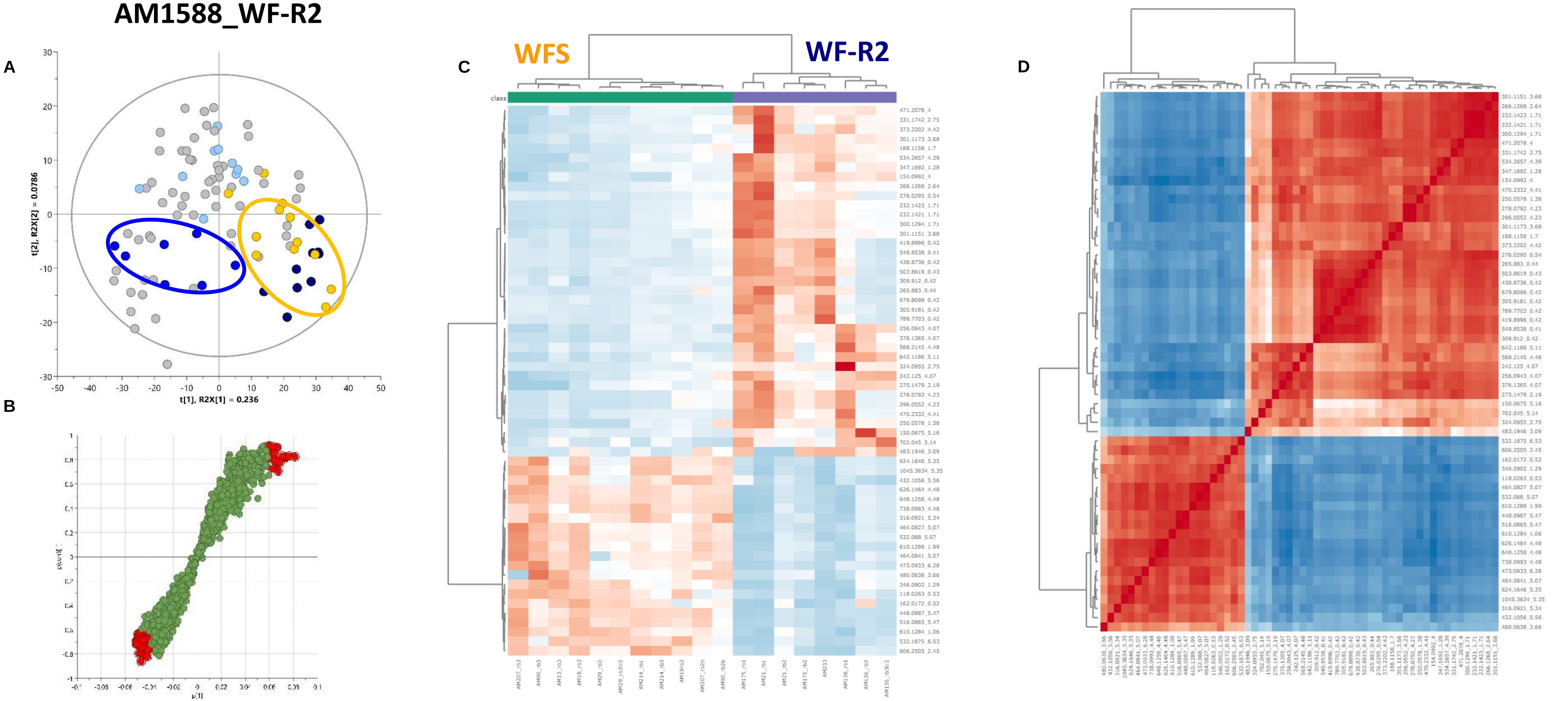

**Supplementary Fig.S4**: Metabolite markers explaining phenotypic classification of F2's WF-R2 subgroup and WF-S group.

### Metabolite markers of extreme metabo/phenotypes

Heatmap & dendrogram

Metabolite-metabolite correlation (Pearson's)

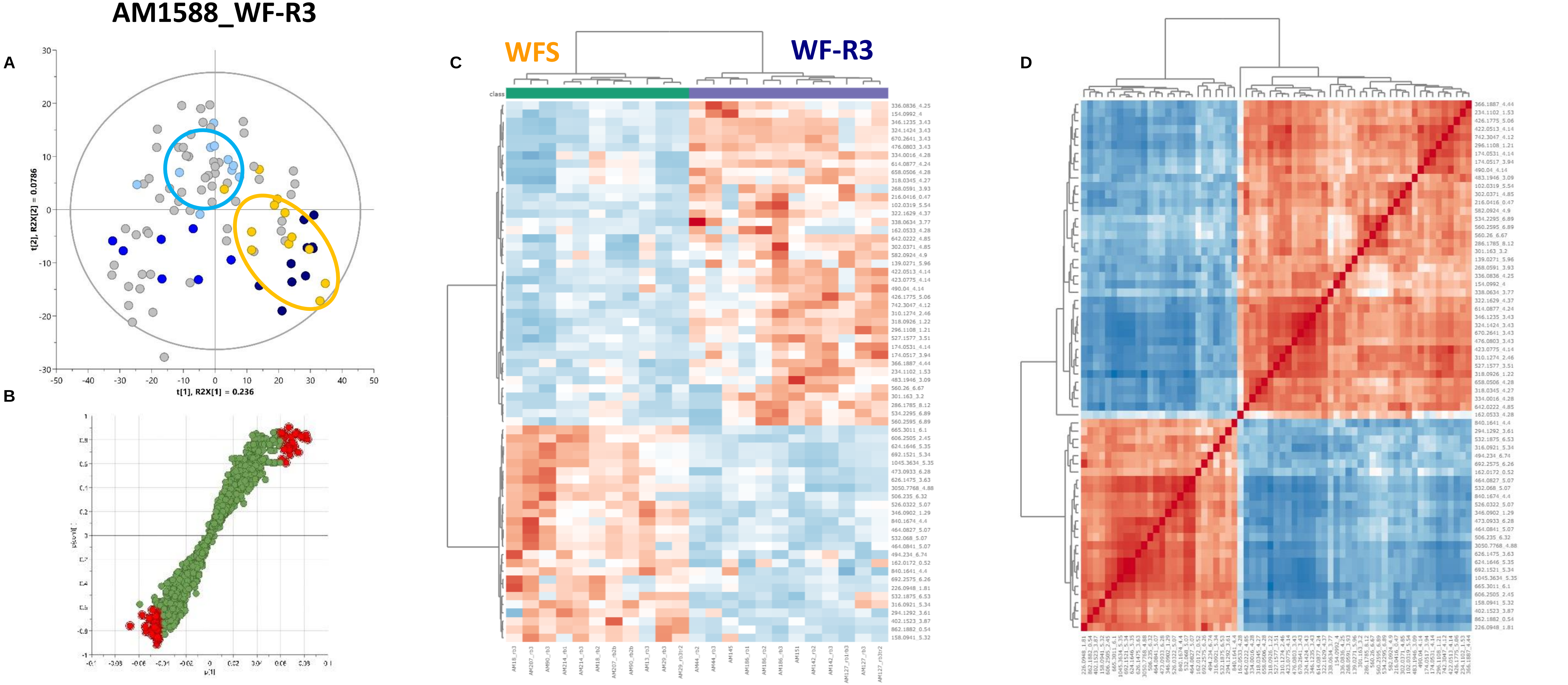

**Supplementary Fig.S5**: Metabolite markers explaining phenotypic classification of F2's WF-R3 subgroup and WF-S group.

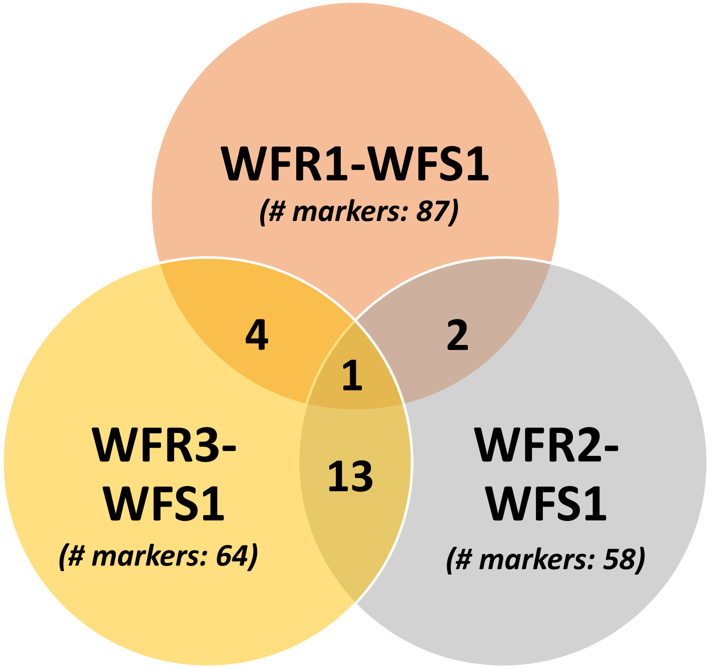

| WFR1-WFR2 | WFR1-WFR3 | WFR2-WFR3 |
|---|---|---|
| 503.8619_0.43 | 402.1523_3.87 | 162.0172_0.52 |
| 316.0921_5.34 | 318.0345_4.27 | 346.0902_1.29 |
| | 658.0506_4.28 | 606.2505_2.45 |
| | 316.0921_5.34 | 483.1946_3.09 |
| | | 154.0992_4 |
| | | 532.068_5.07 |
| | | 464.0827_5.07 |
| | | 464.0841_5.07 |
| | | 316.0921_5.34 |
| | | 1045.3634_5.35 |
| | | 624.1646_5.35 |
| | | 473.0933_6.28 |
| | | 532.1875_6.53 |

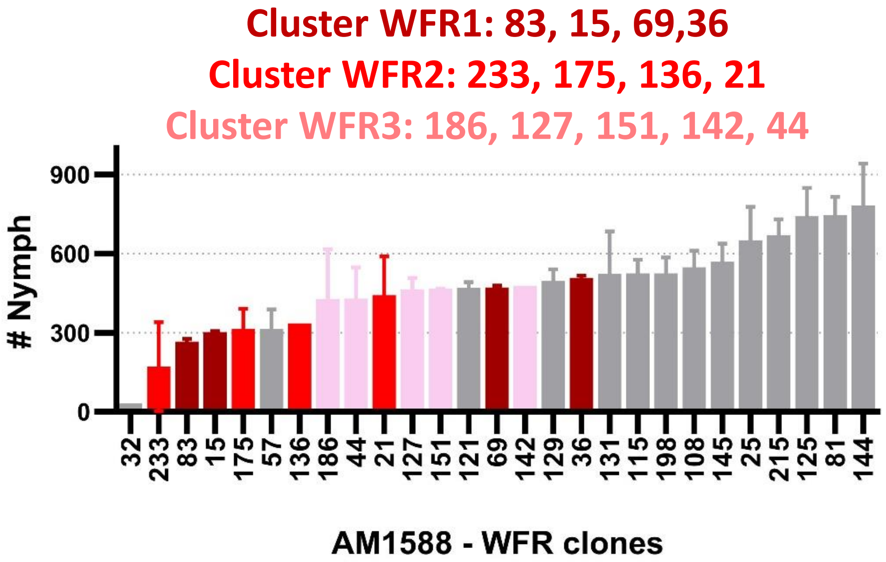

**Supplementary Fig.S6**: Venn diagram of metabolite markers identified within the WF-R class of the F2 family AM1588.
